# Weak interactions cause poor performance of common network inference models

**DOI:** 10.1101/2023.06.22.546059

**Authors:** Dinara Sadykova, Jon M. Yearsley, Andrej Aderhold, Frank Dondelinger, Hannah J. White, Lupe Leon Sanchez, Maja Ilić, Alexander Sadykov, Mark Emmerson, Paul Caplat

## Abstract

1. Network inference models have been widely applied in ecological, genetic and social studies to infer unknown interactions. However, little is known about how well the models perform and whether they produce reliable results when confronted with networks where weak interactions predominate and for different amounts of data. This is an important consideration as empirical interaction strengths are commonly skewed towards weaker interactions, which is especially relevant in ecological networks, and a number of studies suggest the importance of weak interactions for ensuring the dynamic stability of a system.
2. Here we investigate four commonly used network methods (Bayesian Networks, Graphical Gaussian Models, L1-regularised regression with the least absolute shrinkage and selection operator, and Sparse Bayesian Regression) and employ network simulations with different interaction strengths to assess their accuracy and reliability.
3. The results show poor performance, in terms of the ability to discriminate between existing relationships and no relationships, in the presence of weak interactions, for all the selected network inference methods.
4. Our findings suggest that though these models have some promise for network inference with networks that consist of medium or strong interactions and larger amounts of data, data with weak interactions does not provide enough information for the models to reliably identify interactions. Therefore, networks inferred from data of that type should be interpreted with caution.

## Introduction

Network inference models that can infer the “true” structure of networks from data are of great interest and popularity, due to the need to understand the interactions between species, genes, neurones or social groups. Furthermore, it allows the study of subject responses to anthropogenic and non-anthropogenic disturbances, network collapses, and more. For example, epidemiological networks have revealed that the influence of individuals in spreading infections need not be homogeneous, thus informing public health interventions (Dangerfield et al., 2008). Knowledge of gene networks might be useful to assess possible side effects of existing drugs and compounds (Lee et al., 2011). Inferring ecological networks might help to unveil the complexity of some ecosystems and guide conservation (Kaiser-Bunbury et al., 2015).

Depending on the system under investigation, interaction strengths between network nodes (e.g. species, genes, agents) may vary substantially, as well as the size of the network (number of nodes) and the amount of data available. Many studies use simulations to investigate performance of network inference methods, in terms of the ability of the methods to infer the correct network structure (connections between the nodes). Several other studies show that empirically measured interactions strength distributions are generally skewed towards weaker interactions (Berlow et al., 2004; Emmerson and Raffaelli, 2004; Wootton and Emmerson, 2005). However, there is a lack of papers investigating the performance of network inference methods based on nodes with weak, medium or strong interaction strengths for different experimental setups. This lack of performance control is striking for a field with great momentum, as was recently the field of Species Distribution Modelling (Stockwell and Peterson, 2002). Unreliable inference of network structures would have far-reaching consequences in the different areas of application.

In this paper, we fill this gap by employing a simulation approach with different strategies to evaluate performance of four commonly used network inference methods: 1 - Bayesian Networks (BN), 2 - Graphical Gaussian Models (GGM), 3 - L1-regularised regression with the least absolute shrinkage and selection operator (LASSO) and 4 - Sparse Bayesian Regression (SBR). The simulation approach allowed us to vary interaction strength, the number of nodes, the sample size (number of sites/experiments, etc.) and network connectivity. The performance of each method for identifying the presence or absence of a pairwise interaction was evaluated using the Area Under the Receiver Operating Characteristics (AUROC) score (Davis and Goadrich, 2006). We aimed to (1) assess the effectiveness of the four network methods for inferring networks with different average interaction strengths; (2) identify the areas (in terms of network size and sample size) where the methods’ estimates are reliable (reliability is defined in Methods, AUROC scores subsection).

## Materials and Methods

### Terminology

Nodes - species, genes, neurones, groups, or any element of interest. Note that here we focus on ecological interactions but results are generalizable to any type of network.

Edges - connections between the nodes

Sample size - number of locations, sites, samples, experiments, etc.

Interaction strength of node A upon node B is defined here as the change in the growth rate at node B given by a unit increase in node A.

### Simulated Network data

1. Simulation data from ecological simulation model. We used ecological simulation models following Faisal et al. (2010) and Aderhold et al. (2012, 2013) to generate population data. The ecological simulation model is a combination of two models: a trophic niche model (Williams & Martinez, 2000; Faisal et al., 2010; Aderhold et al., 2012, 2013) and a stochastic population model on a 2-dimensional lattice (Lande et al. (chap. 8), 2003; Faisal et al., 2010; Aderhold et al., 2012, 2013). The trophic niche model defined the structure of the network and has only two parameters: the number of species and network connectivity. Each node was randomly given a “niche value”, drawn uniformly from [0; 1] to allocate its place in the food chain (the higher “niche value” it has, the higher up the node is in the food chain; and species are constrained to consume all prey species within their “niche range”). For each node a “niche range” (the size of the niche that the node preys on) has also been determined from a beta distribution with an expected value of 2 multiplied by network connectivity (values of network connectivity are found in Supplementary Materials, Table S1). This determined the size of the niche that the species prey upon. Then a centre for the niche was drawn from the Uniform [0.5^*^growth rate; niche value] distribution. Networks generated from the niche model have been shown to share characteristics with the real food webs (such as existence of a fraction of nodes with or without prey or predator, the amount of cannibalism, etc.). It should be noted here that the classic formulation of a niche space, as a space corresponding to ecological and environmental characteristics, is not applicable here. This model is restricted to a species feeding niche and assembles hierarchy of interactions among species (Williams and Martinez, 2000). The population model was defined by a stochastic differential equation:

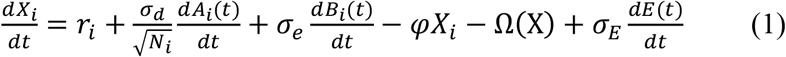

Where the dynamics of log abundance *X*_*i*_ of one species *i* was a function of its growth rate *r*_*i*_, the species-specific demographic effect *A*_*i*_*(t)* (where *N*_*i*_ is the abundance of species 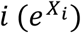, the standard deviation of the demographic effect σ_*d*_, the species-specific environmental effect *B*_*i*_(*t*), the standard deviation of the species-specific environmental effect σ_*e*_, the intra specific density dependence *φ*, effect of competition for common resources Ω, the general community environment *E*(*t*) and the standard deviation of the general environmental effect σ_*E*_ (following Faisal et al., 2010). In order to incorporate the niche model described above to the population model (1), we modified the term Ω (i.e., competition for resources) when simulating the data, to include predator-prey interactions in the Lotka-Volterra form. All model terms (i.e., competition for resources, which depends on the niche values and the niche feeding space, species-specific demographic/environmental effects, etc.) influence population abundance, but otherwise independent. For more details about the model, see Faisal et al. (2010) and Aderhold et al. (2012, 2013). Further, the simulation employed an exponential dispersal model to extend the model to a 2D arena. Here the probability of a species moving from location A to location B was determined by the Euclidean distance between A and B. Each location had its own growth rate and spatial pattern of growth rates for one species was generated by noise with spectral density *f*^*β*^ (*β <* 0, *f* is the frequency at which the noise is measured), and a normal error distribution. We considered sample sizes of 25, 100, 225, 400 and 625 (based on rectangular grids from 5x5 to 25x25 with a step equal to 5) and the number of nodes of 5, 10 up to 150 with a step equal to 10 starting from 10 species (for weak interaction models) and from 5 to 50 with the same step as before (for medium/strong interaction models; due to the high number of extinct nodes when medium/strong interactions were used, it was technically very difficult to simulate a large number of nodes) (Table S1^1^). For each node-sample size pair, we selected the mean non-zero network interaction strength to be 1, 2 or 4 (Fig. S9^1^), where 1 stands for “weak” interactions, 2 - for “medium” and 4 - for “strong” interactions. The networks with any type of interaction strengths were mainly filled with zeros, with relatively few non-zero interactions, which are shown in Fig S9. In this study, we also selected network connectivity (network density) from 0.05, 0.1 or 0.15 (all the main results are given for a connectivity of 0.1). All other model parameters (intra-specific density-dependence, variation in growth rate and spatial autocovariance) were constant across all the simulations (Table S1). The dynamics of this model were simulated for 3000 steps until the system had reached the equilibrium. The simulations were performed 200 times for each node-sample size pair, for different network interaction strength values and network connectivity values. The simulation data consists of species abundance data (the considered number of species (i.e., nodes) and the sample sizes are provided above), which are used to reconstruct the interaction network. The known interaction network (a matrix with ones and zeros for presence/absence of interactions between two species) is also provided, which is assumed to be unknown during the reconstruction process.
2. Real food webs. Food webs from seven locations: (a) Baltic Sea; (b) Broadstone stream; (c) Lake Vättern; (d) Montane forest; (e) Skipwith Pond; (f) Tropical Sea and (g) Trelease Woods were incorporated into the ecological simulation models (one food web was considered at a time) to understand the distribution of interaction strengths for food webs and what results would be obtained for the food webs based on empirical topology and the observed body mass spectrum. The interaction strengths were taken from Berg et al. (2011), Woodward et al. (2005), Cohen et al. (1990), Warren (1999), Brose et al. (2005), Säterberg et al. (2019) and we implemented 200 replicates for each food web.
3. Simulation data from Graphical Gaussian Models (GGM). 200 random networks (which are based on the partial correlation matrices in GGM) were generated for each node-sample size pair, and data were simulated from the network-corresponding multivariate normal distribution using “GeneNet” library in R (Schafer and Strimmer, 2005). We considered 1000 and 3000 nodes and 100, 500 and 1000 sample sizes to understand whether models are more reliable for a large number of nodes (say, gene studies).

### Methods

1. Bayesian Networks (BN). BN are probabilistic graphical models, which means that they can define relationships between nodes and can be used to calculate probabilities that define the chance of finding a node in a given state (by “state” we mean presence or absence of an interaction) (Grzegorczyk and Husmeier, 2008; Faisal et al., 2010; Aderhold et al., 2012, 2013). BN is defined by a graph, which consists of a set of nodes and a set of edges between the nodes. Each node corresponds to a unique random variable and the edges represent interactions between them. The BNs interaction network was considered here as an undirected acyclic graph, where “acyclic” means that no one node can interact with themselves and “undirected” means that the detected relationships could indicate either one or two way relationships, without a direction of influence. For further details, see Grzegorczyk and Husmeier (2008), Faisal et al. (2010) and Aderhold et al. (2012, 2013). We employed Markov Chain Monte Carlo (MCMC) learning methods with a special edge reversal move by Grzegorczyk and Husmeier (2008). The MCMC method started a chain with an empty graph and it was either adding, deleting or reversing an edge at each step. The algorithm was further extended to produce a network that can deal with spatial autocorrelation (when nearby locations are more similar than the locations that are further away). For this reason, the average population at neighbouring cells was computed and weighted inversely proportional to the Euclidean distance of the neighbours, which was then defined as an autocorrelation variable. Each node was then considered to be linked to a node that represents the autocorrelation. In this way, the observation status of a node was firstly predicted by the spatial neighbourhood. BN modelling was performed in MatLAB (https://github.com/FrankD/EcoNets).
2. Graphical Gaussian Model (GGM). GGM is a graphical model that is represented by an undirected network of partial correlation coefficients. The GGM assumes multivariate Gaussian distribution of the data and estimates the inverse covariance matrix of this distribution to obtain partial correlation coefficients and then the conditional independence graph (conditional on all the other nodes of the given network). The partial correlations are graphically displayed as a weighted network with nodes and edges between them, which represent the partial correlation between two nodes. When the partial correlation is zero, the edge is not present. For more details, see Lauritzen (1996, 2004), Edwards (2000) and Faisal et al. (2010). GGM modelling was performed using the ggm.estimate.pcor function in R (GeneNet package: Graphical Gaussian Models: Small Sample Estimation of Partial Correlation).
3. Least absolute shrinkage and selection operator (LASSO). LASSO is an approach that performs estimation, variable selection and regularization in regression models (Tibshirani, 1996; Faisal et al., 2010). LASSO regression model is a way to model relationships between a given target variable and other dependent variables. These relationships are modelled with linear coefficients (also named as regression weights). LASSO regression shrinks and regularizes these regression coefficients (weights) using l1-penalization, where penalty depends on a tuning parameter *λ* that controls the amount of shrinkage, to improve prediction accuracy. This shrinkage drives coefficients to zero by penalising large regression coefficients. We interpret the coefficients as edge weights in the network. The regularisation (hyper-) parameter defines the sparsity of the inferred network. The tuning parameter is chosen using cross validation. When *λ* is small, the results are similar to the least squares estimates, as *λ* increases, more coefficients are driven to zero and thrown away. In this paper we used the tuning parameter that provided minimum mean cross-validated error. We also tested other tuning parameters, but they provided worse results of network restoring. The LASSO interaction network was considered as an undirected graph. LASSO modelling was performed using the cv.glmnet function in R (glmnet package: Lasso and Elastic-Net Regularized Linear Models) with 10 folds cross validation.
4. Sparse Bayesian Regression (SBR). SBR is a method, which employs a sparse regression method with a probabilistic model (Tippin, 2001; Rogers and Girolami, 2005; Faisal et al., 2010). By selecting a small number of non-zero effects from a large number of candidate effects, SBR method produces networks that are naturally sparse. Similar to the LASSO method, the SBR method uses a tuning parameter. The SBR models relationships between a node and other nodes in a regression model in order to find edges between them. The found set of nodes with edges between the given node and this set of nodes, is produced with coefficients (weights), which show interaction strength between them. The interaction strength matrix is developed based on the weight matrix and the network is defined as a set of interactions with non-zero interaction strength. The SBR interaction network was considered as an undirected graph. SBR modelling was performed using sparsereg function in R (sparsereg package: Sparse regression for experimental and observational data).
5. Data for the methods. All the methods (BN, GGM, LASSO and SBR) reconstruct the interaction networks using the species abundance data, produced by the simulations. The reconstructed networks are produced as either undirected acyclic graph scores (BN), absolute values of sparse estimates (LASSO, SBR) or estimators of partial correlation (GGM). The reconstructed networks are then compared to the true network, which are known precisely, using AUROC scores.
6. Interaction strength. Each network inference model has a different definition of interaction strengths: GGM considers them as partial correlation coefficients, LASSO and SBR as regularised regression coefficients and BN as marginal posterior probabilities. The common thing between these definitions is that they all define a ranking of the edges, which can be used to estimate AUROC scores (Faisal et al., 2010).
7. The AUROC scores are calculated based on a receiver operating characteristic (ROC) curve, where the true positive rate (relative number of true interactions) is plotted against the false positive rate (relative number of false interactions) for all possible thresholds on the rank (Davis, 2006). AUROC ranges in value from 0 to 1, where zero shows that all predictions are 100 percent wrong, while one shows that all predictions are 100 percent accurate. When AUROC score is 0.5, the model has no discrimination capacity to distinguish between present and absent interactions. AUROC scores greater than 0.7 are considered to be acceptable for good predictions in this manuscript; scores greater than 0.8 would show excellent performance and greater than 0.9 - outstanding performance of a model (Mandrekar, 2010). Ideally, for simulation studies, the percentage of the AUROC scores in the range between 0.7 and 1.0 should be 100% to demonstrate that a model shows only acceptable scores for any simulated data. However, it was not always the case and therefore, in this study, we introduced two categories: (1) reliable models - models that showed 98% or more of the acceptable AUROC scores. This means that among 200 simulation runs the model showed AUROC scores less than 0.7 for only 4 simulations (2%) or less and the model performs network inference with 98% reliability; (2) non-reliable models - models that showed 50% or less of the acceptable AUROC scores. This means that the model showed unacceptable AUROC scores for every second simulation or more. All other models that are uncategorized, will be referred as “less-reliable models” with some degree of reliability, say, a model with 85% would be referred in this paper as a model with 85% reliability. A set of models, say, between 75% and 97%, would be referred to as models with 75%-97% of reliability.

## Results

All four network methods had poor performance at estimating a network’s topology when weak interaction strengths were considered (Fig. 1, 2). Performance was average for interactions with medium strength (Fig. S1, S3) and good for strong interactions (Fig. S2, S4) (interaction strength densities are found in Fig. S9).

**Fig. 1.**
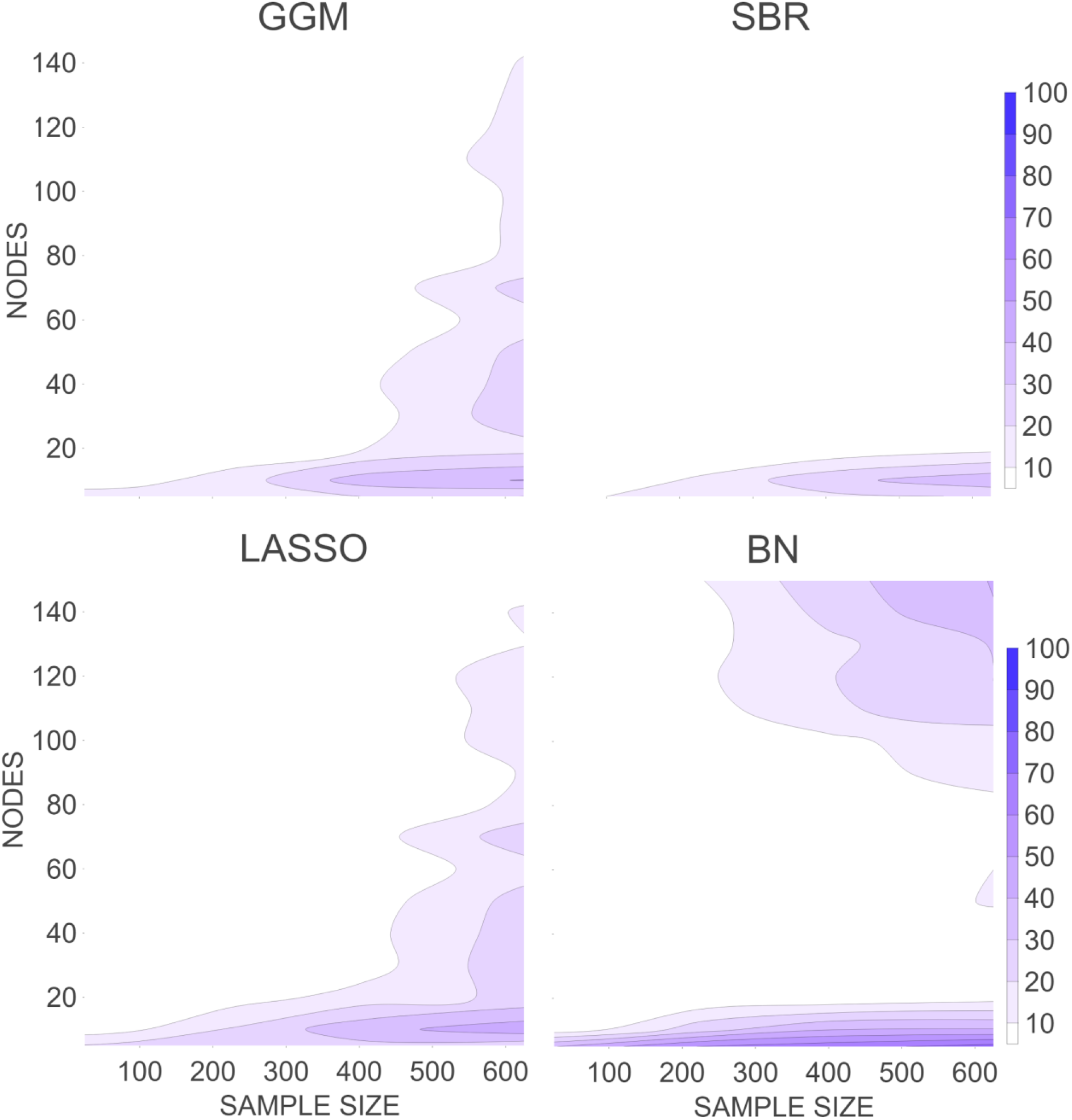
Percentage of the AUROC scores in the range between 0.7 and 1.0, which are considered acceptable scores, for four different methods (GGM - top left; SBR - top right; LASSO - bottom left; BN - bottom right) for simulations with weak interactions. The results are given for different numbers of simulated nodes (y-axis) and sample sizes (x-axis). Dark purple/blue areas show higher percentages of acceptable scores and therefore more reliable models, while white/light purple areas reveal non reliable models.

**Fig. 2.**
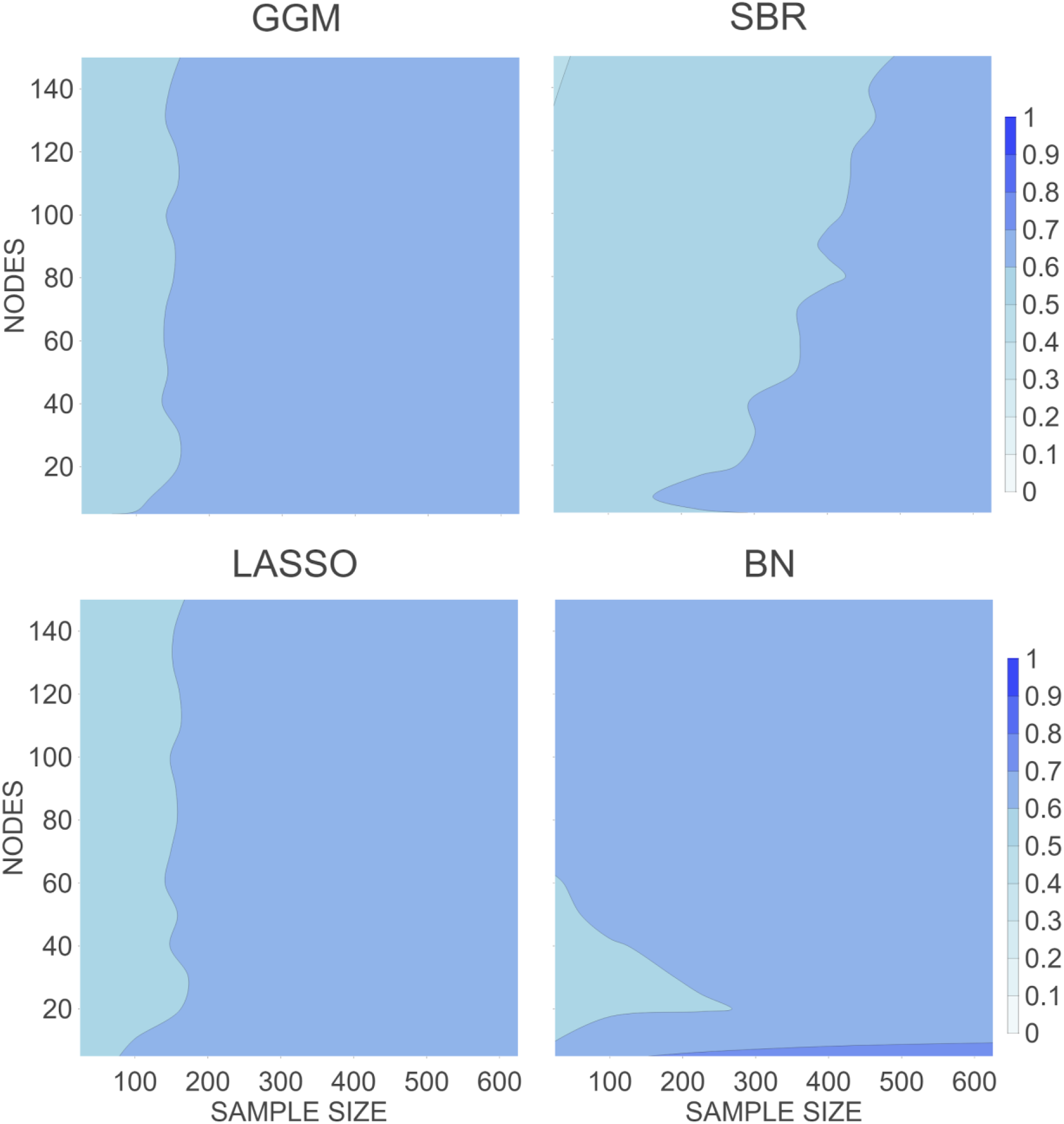
Mean AUROC scores for four different methods (GGM - top left; SBR - top right; LASSO - bottom left; BN - bottom right) for simulations with weak interactions. The results are given for different numbers of simulated nodes (y-axis) and sample sizes (x-axis).

The inference performance for weak interaction networks was substantially lower in comparison to networks with medium or strong interactions (Fig. 1, Fig. S2-S3, Table S2). According to our AUROC scores classification, GGM, LASSO and SBR models did not produce reliable results for all combinations of nodes-sample sizes for weak interactions (Table S2, Fig.1). BN also only gave reliable network inference for networks with 5 nodes and larger sample sizes (225-625 samples).

Performance of the inference models for inferring networks with medium interaction strengths (Fig. S1, S3) identified that a sample size equal to 625 with a network containing at least 30 nodes gave reliable inference for all four methods. In addition, sample sizes greater than 400 with networks containing at least 20 nodes produced reliable network inference for all methods except BN (which showed 94-95% reliability). LASSO and GGM also showed that a sample size equal to 225 with a network containing at least 40 nodes gave reliable inference. At the same time, a sample size of 25 did not give reliable inference for all the models.

Results of performance of the inference models for inferring networks with medium interactions indicated that all four methods performed well, producing reliable results for ≥100 sample sizes with ≥10 nodes (Fig. S3), except a combination of 10 nodes with the sample size = 100 for GGM, SBR and LASSO methods, which showed slightly smaller reliability (97% - GGM, 96% - SBR and 90% - LASSO). When inferring networks with strong interactions, all of the models were reliable, except for the SBR method, which showed non-reliable results with the sample size = 25 and ≥30 nodes. Results of the AUROC scores for different network models are found in Fig. 3, Fig. S5-S7.

**Fig. 3.**
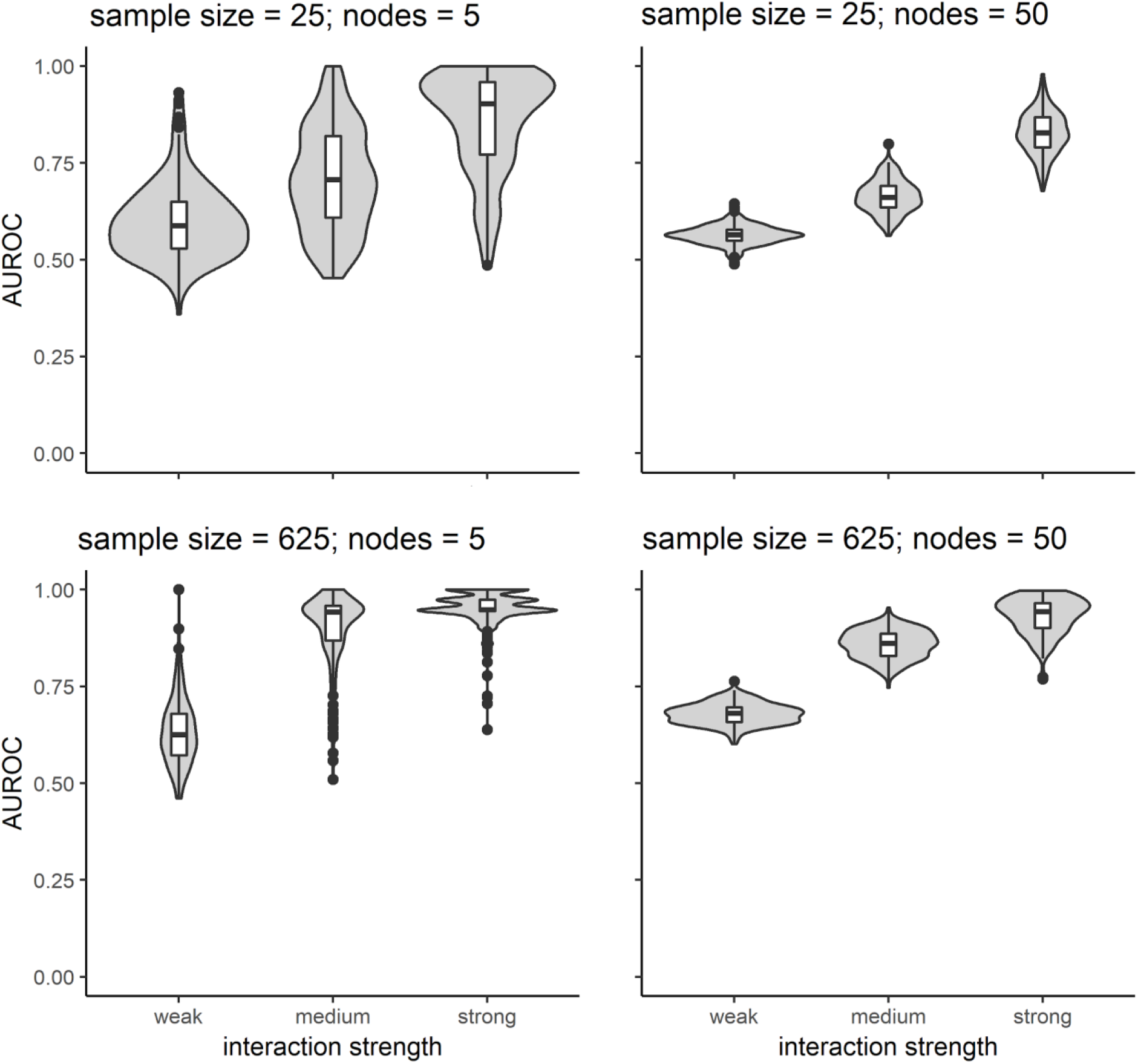
The AUROC scores for the GGM method. The plots show kernel probability density of the scores for three levels of interaction strengths (weak - left; medium - middle; strong - right) and different sample sizes and numbers of nodes (sample size equals to 25 - top; sample size equals to 625 - bottom; 5 nodes - left; 50 nodes - right). Box plots that are included inside the violin plots are showing median values, the 25th and 75th percentiles and minimum/maximum values of the AUROC scores within distance equal to interquartile range multiplied by 1.5. The dots show the points that are beyond that distance.

Results based on the GGM simulated data with 1000 and 3000 nodes and sample sizes of 100, 500 and 1000 showed that GGM performance does not improve as more nodes are considered. In fact, all the obtained AUROC scores were below 0.58 showing that the models had no discrimination capacity to distinguish between present and absent interactions.

Many published studies fall in a parameter space (in terms of number of nodes/sample size) that we found would lead to non-reliable results with networks that are made with weak or medium interaction strengths (Fig S10). Out of 25 ecological studies that used network inference models, 68% were inside the considered parameter space. Among these, 33% and 100% were inside the non-reliable areas when considering networks with medium or weak interaction strengths, respectively, for all the methods studied here.

Results of performance of the inference models can also be analysed by mean (average) AUROC scores (Fig 2, S3 and S4), which indicate average model performance. The AUROC mean values for GGM, LASSO and SBR models varied between 0.55 and 0.69 scores for models with weak interactions, indicating that, on average, the models had no discrimination capacity to distinguish between present and absent interactions. The AUROC mean values for BN models varied from 0.56 to 0.75 scores for models with weak interactions, showing that only combinations of 5 nodes with sample sizes 225-625 or combinations of 140-150 nodes with the sample size of 625 would show acceptable scores on average.

The considered methods varied in their performance (Fig. 1-2, Fig. S1-S2), with the SBR method showing significantly lower AUROC values than other network methods for simulations with the largest considered number of sample sizes (625) when the number of nodes was high (50) (for weak interactions) or with a smaller sample size (25), with a number of nodes 20-50 (for medium and strong interactions) (Table S2, Fig. 1-2, S1-S4). For example, the SBR method had a reliability of 0%, whilst the BN method had a reliability of 40.5%, when applied to a network containing 150 nodes with weak interactions and a sample size of 625. For networks with strong interactions, 50 nodes and sample size of 25, the SBR method showed 9.5% reliability (i.e., non-reliable), while GGM, LASSO and BN showed 99% (almost perfect reliability), 76.5% and 85.5% reliability respectively.

BN performed better than the other methods when networks made of weak interactions were considered (Table S2, Fig. 1-2). However, for medium interactions, even though the BN method performed better than other methods for a small number of nodes (<10), BN slightly underperformed for a larger number of nodes (≥20) with a large sample size (≥400).

Performance of the network inference models with weak interaction strength and different network connectivity values (Fig. S8) showed that network connectivity played a minor role in estimating a models’ performance and the estimated AUROC scores were similar for most of the combinations of node-sample size for different connectivity values. In particular, the results were nearly identical for a large number of nodes (≥20) with any sample size, and slightly better results for 5-10 nodes, 625 sample size and smaller network connectivity (0.05) (but the estimated reliability was only a bit outside of the non-reliable region).

Overall, the results show steep drops in performance of all the methods for all interaction strengths when the sample size is small (<300).

This study allowed us to identify areas (in terms of number of nodes, sample size and interaction strengths) where the methods’ estimates are reliable, which can be seen in Fig. S1- S2 (Supplementary Materials).

## Discussion

Our study shows that, with a large sample size, and medium or strong interactions, inference network models perform well. However, weak interactions cause a strong drop in model performance, which puts such studies into questions. Ecological networks commonly have relatively weak interactions that are essential in the functioning of these networks. We, therefore, encourage authors to look for a priori knowledge about the nodes’ interactions before using these methods and consider using a subset of stronger connected nodes.

We compared the results based on simulation models described above with the results based on food webs with empirical topology. We found that the food web outcomes correspond to the outcomes of the simulation models with weak interaction strength, where non-reliable results were produced for all node-sample size pairs. This finding suggests that data with weak interactions (thought to be prevalent in ecological webs and other networks (Berlow et al., 2004; Emmerson and Raffaelli, 2004; Wootton and Emmerson, 2005) do not provide enough information for the inference network models to reliably identify interactions using the four methods studied here unless network size is small and sample size is large.

It should be acknowledged here that the measure of interaction strengths might vary for different studies (Berlow et al., 2004). Berlow et al. (2004) showed that there are several different definitions of the term “interaction strengths” and they measure different properties, for example, elements of the community matrix, statistical correlations among abundances or relative prey preferences. Therefore, the results from this paper must be interpreted in the context of our definition of interaction strength of node A upon node B as the change in the growth rate at node B given by a unit increase in node A.

Here, we would like to caution that methods with regularisation such as SBR or LASSO might estimate small/weak effects poorly as small effects are biased towards zero by these methods. We employed these two methods here to show that some of the common network methods are possibly invalid for networks with weak interactions, while researchers keep using these methods. We would also like to note that the tuning parameters were selected to show the best LASSO and SBR results (taking into account the model parameters, Table S1, Supplementary Materials), while other tuning parameters produced either similar or worse results than those we are publishing. This means that tuning parameters that produce less shrinkage up to almost no shrinkage add more regression coefficients (network connections), which are mismatched with the true connections and therefore AUROC gets smaller. While more shrinkage removes some of the detected matched connections.

However, it should be noted that we only considered network connectivity 0.05-0.15 in this ms, based on the real food webs (Table S1, Supplementary Materials). If network connectivity equals to one (i.e., all nodes are connected), then selecting a tuning parameter with no shrinkage will always provide perfect AUROC scores. Therefore, network connectivity should be taken into account when selecting a tuning parameter.

This paper will help researchers to understand that network inference models are appropriate to use when the strength of interactions is either medium or strong, with sufficiently large sample and node sizes, and provides researchers with information about experimental conditions under which these network methods are accurate and consistent.

## Acknowledgments

Willson Gaul provided comments on a draft of the manuscript.

## Author contributions

Dinara Sadykova contributed to the conception and design of the work, contributed to the analysis and interpretation of the data for the work; drafted and revised the article; approved the final version to be published;

Jon Yearsley contributed to the conception and design of the work; contributed to the analysis and interpretation of the data for the work; drafted and revised the article; approved the final version to be published;

Andrej Aderhold contributed to the analysis of the work; revised drafts of the paper, approved the final draft.

Frank Dondelinger contributed to the analysis of the work; revised drafts of the paper, approved the final draft.

Hannah White drafted and revised the paper, approved the final draft. Lupe Leon Sanchez revised the paper, approved the final draft.

Maja Ilić drafted and revised the paper, approved the final draft.

Alexander Sadykov contributed to conceptualization of the paper; drafted and revised the paper, approved the final draft.

Mark Emmerson contributed to the conception and design of the work; contributed to the analysis and interpretation of the data for the work; revised the article; approved the final version to be published;

Paul Caplat contributed to the conception and design of the work; contributed to the analysis and interpretation of the data for the work; drafted and revised the article; approved the final version to be published;

## Data Statement

This is a simulation study and the code to produce and analyse the data is found here: https://github.com/FrankD/EcoNets. This manuscript does not contain any data to be archived.

## Footnotes

1 Tables S and Figures S are found in Supplementary Materials

