## Supplementary Materials for "Weak interactions cause poor performance of common network inference models"

Table S1. Model parameters

| | |
|---|---|
| Sample size | 25, 100, 225, 400, 625 |
| Number of nodes | 5, 10, 20, 30, 40, 50, 60, 70, 80, 90, 100, 110, 120, 130, 140, 150 |
| Interaction strength | 1 = weak, 2=medium, 4=strong |
| Network connectivity | 0.05, 0.1, 0.15 |
| Intra-specific density-dependence | 5 |
| Variation in growth rate | 5 |

Table S2. The performance of the four methods on networks with weak interaction strengths with varying sample size and nodes. The numbers represent the rounded percentages of the AUROC scores in the range between 0.7 and 1.0. Models with scores in this range are considered reliable.

| nodes | GGM | | | | | SBR | | | | | LASSO | | | | | BN | | | | |
|---|---|---|---|---|---|---|---|---|---|---|---|---|---|---|---|---|---|---|---|---|
| | sample size | | | | | sample size | | | | | sample size | | | | | sample size | | | | |
| | **25** | **100** | **225** | **400** | **625** | **25** | **100** | **225** | **400** | **625** | **25** | **100** | **225** | **400** | **625** | **25** | **100** | **225** | **400** | **625** |
| **5** | 15 | 15 | 16 | 20 | 20 | 8 | 10 | 15 | 18 | 21 | 21 | 24 | 26 | 27 | 24 | 40 | 48 | 61 | 66 | 75 |
| **10** | 5 | 7 | 15 | 35 | 41 | 2 | 5 | 12 | 27 | 37 | 5 | 9 | 21 | 37 | 46 | 6 | 11 | 27 | 41 | 42 |
| **20** | 2 | 0 | 2 | 10 | 16 | 0 | 1 | 1 | 1 | 7 | 1 | 2 | 5 | 14 | 23 | 1 | 3 | 2 | 3 | 7 |
| **30** | 0 | 0 | 1 | 5 | 27 | 0 | 0 | 0 | 1 | 9 | 0 | 0 | 1 | 5 | 28 | 0 | 0 | 1 | 2 | 4 |
| **40** | 0 | 0 | 1 | 8 | 24 | 0 | 0 | 0 | 1 | 3 | 0 | 0 | 2 | 7 | 25 | 0 | 0 | 1 | 1 | 5 |
| **50** | 0 | 0 | 1 | 5 | 23 | 0 | 0 | 0 | 0 | 2 | 0 | 0 | 0 | 5 | 24 | 0 | 0 | 1 | 2 | 11 |
| **60** | 0 | 0 | 0 | 1 | 16 | 0 | 0 | 0 | 0 | 1 | 0 | 0 | 0 | 2 | 16 | 0 | 0 | 3 | 4 | 10 |
| **70** | 0 | 0 | 0 | 3 | 24 | 0 | 0 | 0 | 0 | 1 | 0 | 0 | 0 | 5 | 26 | 0 | 0 | 4 | 6 | 10 |
| **80** | 0 | 0 | 0 | 0 | 12 | 0 | 0 | 0 | 0 | 1 | 0 | 0 | 1 | 1 | 13 | 0 | 0 | 3 | 7 | 7 |
| **90** | 0 | 0 | 0 | 1 | 12 | 0 | 0 | 0 | 0 | 0 | 0 | 0 | 0 | 1 | 11 | 0 | 1 | 3 | 6 | 14 |
| **100** | 0 | 0 | 0 | 1 | 12 | 0 | 0 | 0 | 0 | 0 | 0 | 0 | 0 | 0 | 16 | 0 | 1 | 5 | 7 | 18 |
| **110** | 0 | 0 | 0 | 1 | 15 | 0 | 0 | 0 | 0 | 0 | 0 | 0 | 0 | 2 | 14 | 0 | 2 | 6 | 20 | 22 |
| **120** | 0 | 0 | 0 | 1 | 13 | 0 | 0 | 0 | 0 | 1 | 0 | 0 | 0 | 0 | 17 | 0 | 2 | 9 | 20 | 31 |
| **130** | 0 | 0 | 0 | 0 | 12 | 0 | 0 | 0 | 0 | 0 | 0 | 0 | 0 | 1 | 10 | 0 | 2 | 8 | 18 | 31 |
| **140** | 0 | 0 | 0 | 0 | 11 | 0 | 0 | 0 | 0 | 0 | 0 | 0 | 0 | 1 | 11 | 1 | 2 | 6 | 23 | 40 |
| **150** | 0 | 0 | 0 | 0 | 8 | 0 | 0 | 0 | 0 | 0 | 0 | 0 | 0 | 1 | 6 | 1 | 2 | 10 | 27 | 41 |

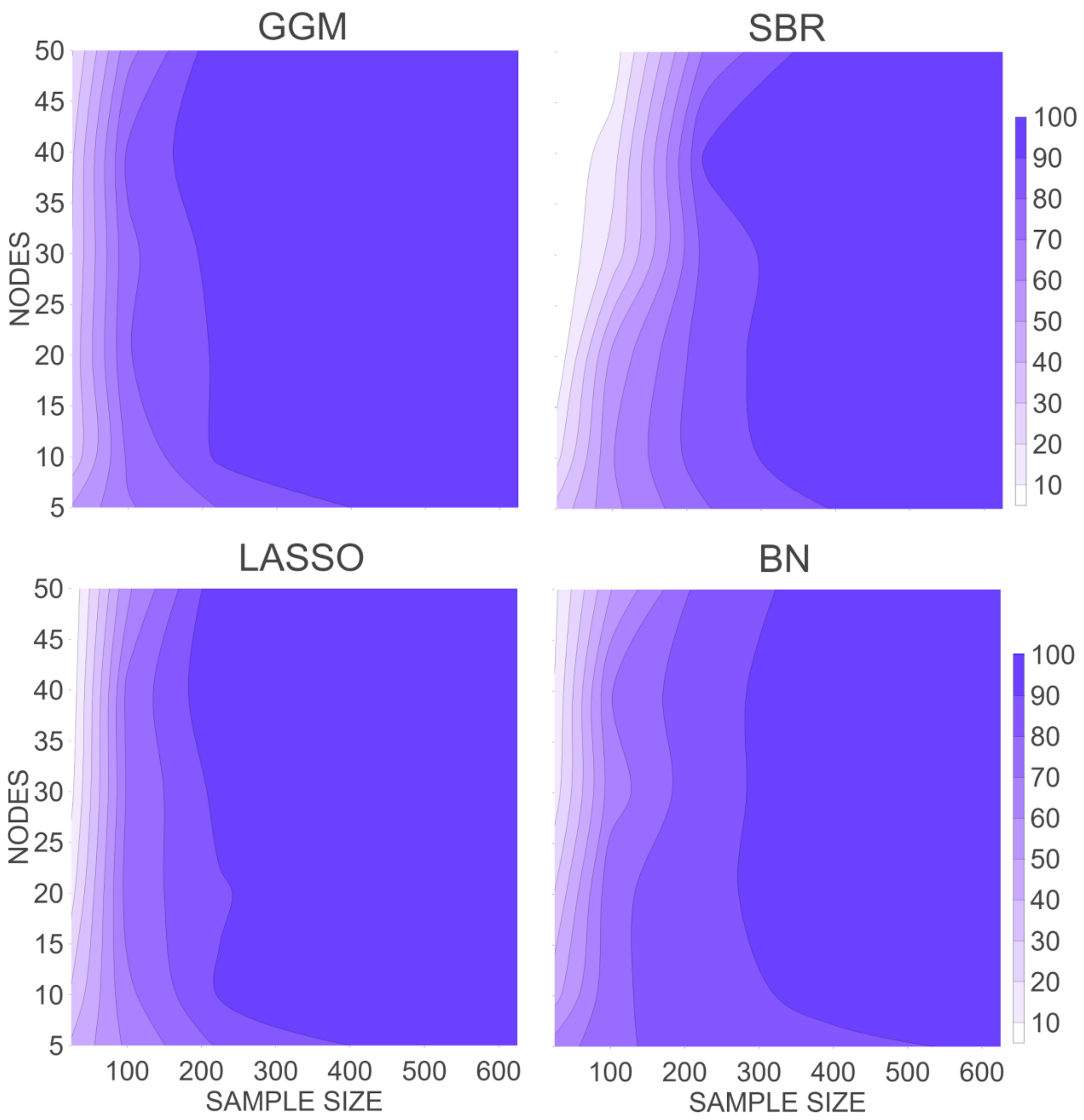


**Fig. S1.**

The performance of the four methods (GGM – top left; SBR – top right; LASSO – bottom left; BN – bottom right) on networks with medium interaction strengths with varying sample size and nodes. The contours and color scale represent the percentage of the AUROC scores in the range between 0.7 and 1.0

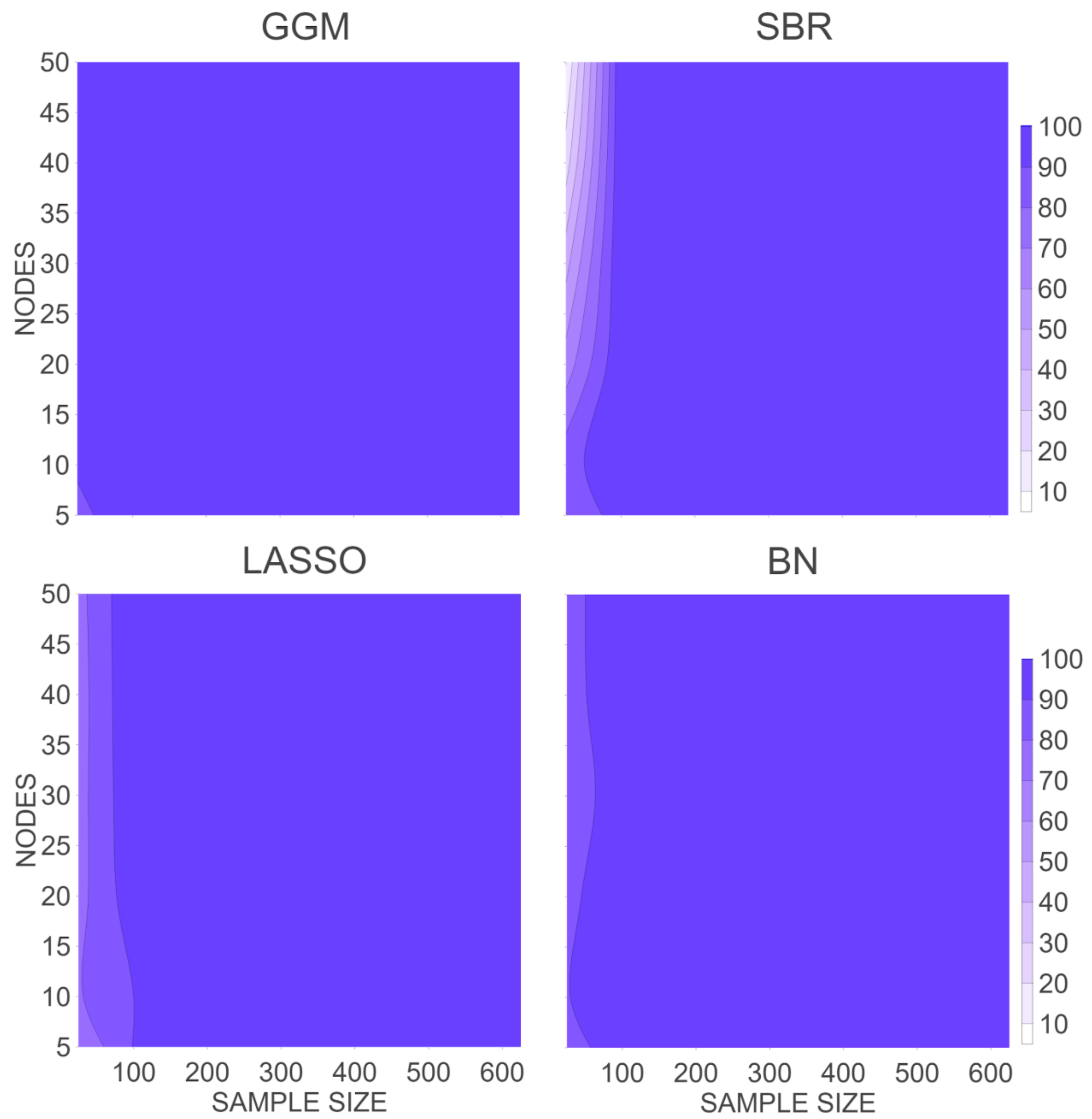


**Fig. S2.**

The performance of the four methods on networks with strong interaction strengths. Color scale and labels are the same as Fig. S1.

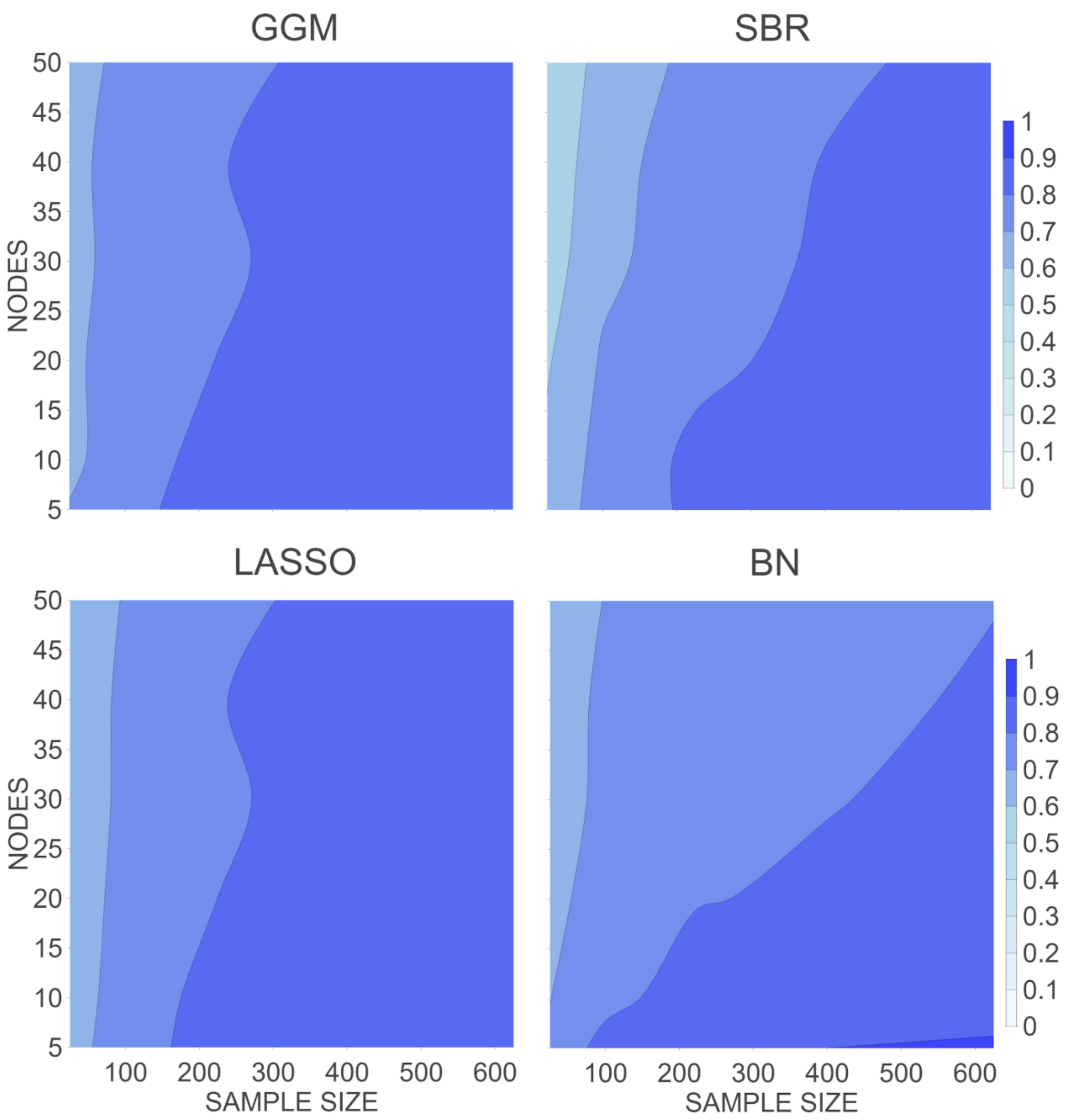


**Fig. S3.**

Mean AUROC scores for the four methods (GGM – top left; SBR – top right; LASSO – bottom left; BN – bottom right) on networks with medium interaction strengths.

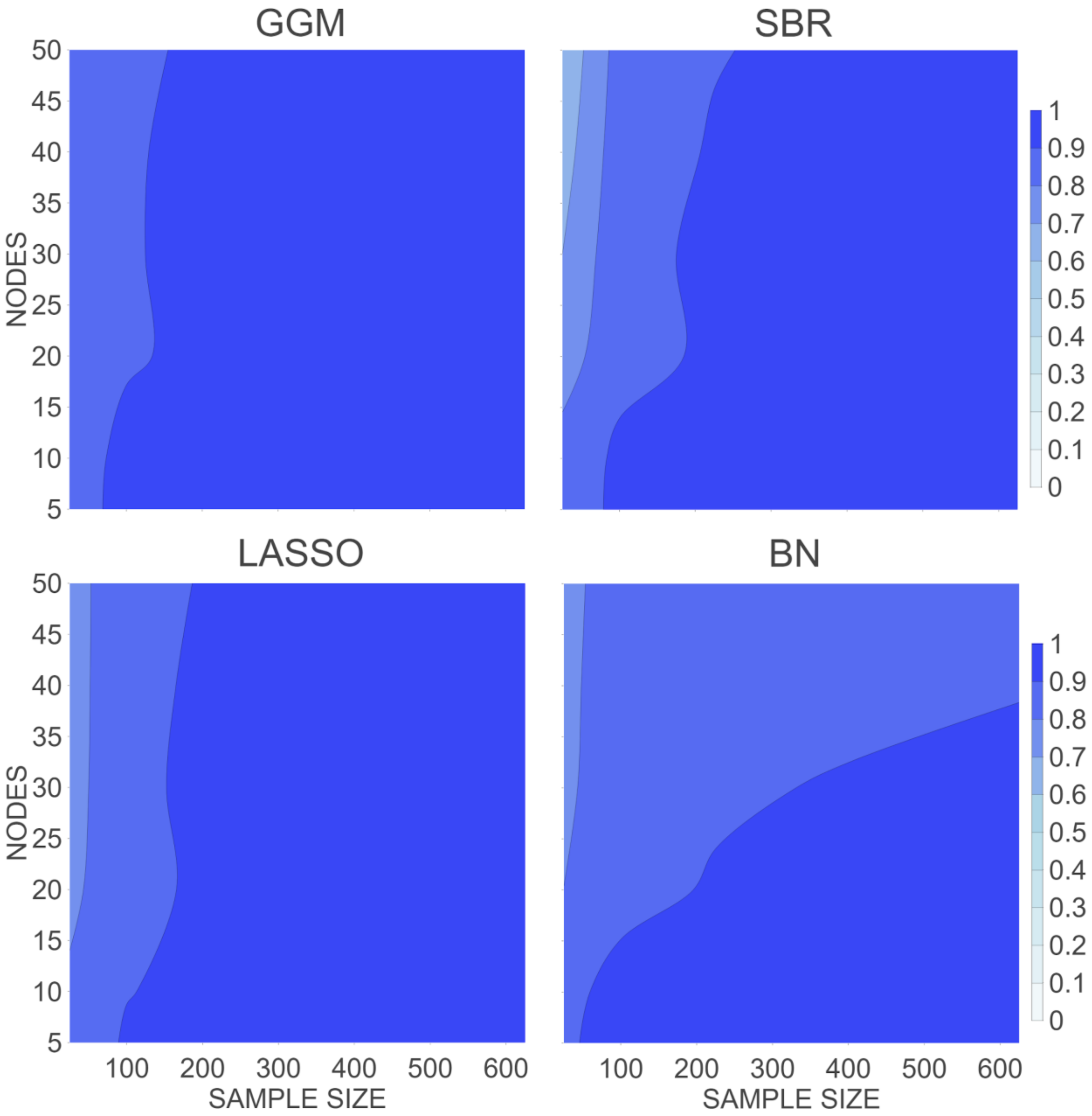


**Fig. S4.**

Mean AUROC scores for the four methods (GGM – top left; SBR – top right; LASSO – bottom left; BN – bottom right), on networks with strong interaction strengths.

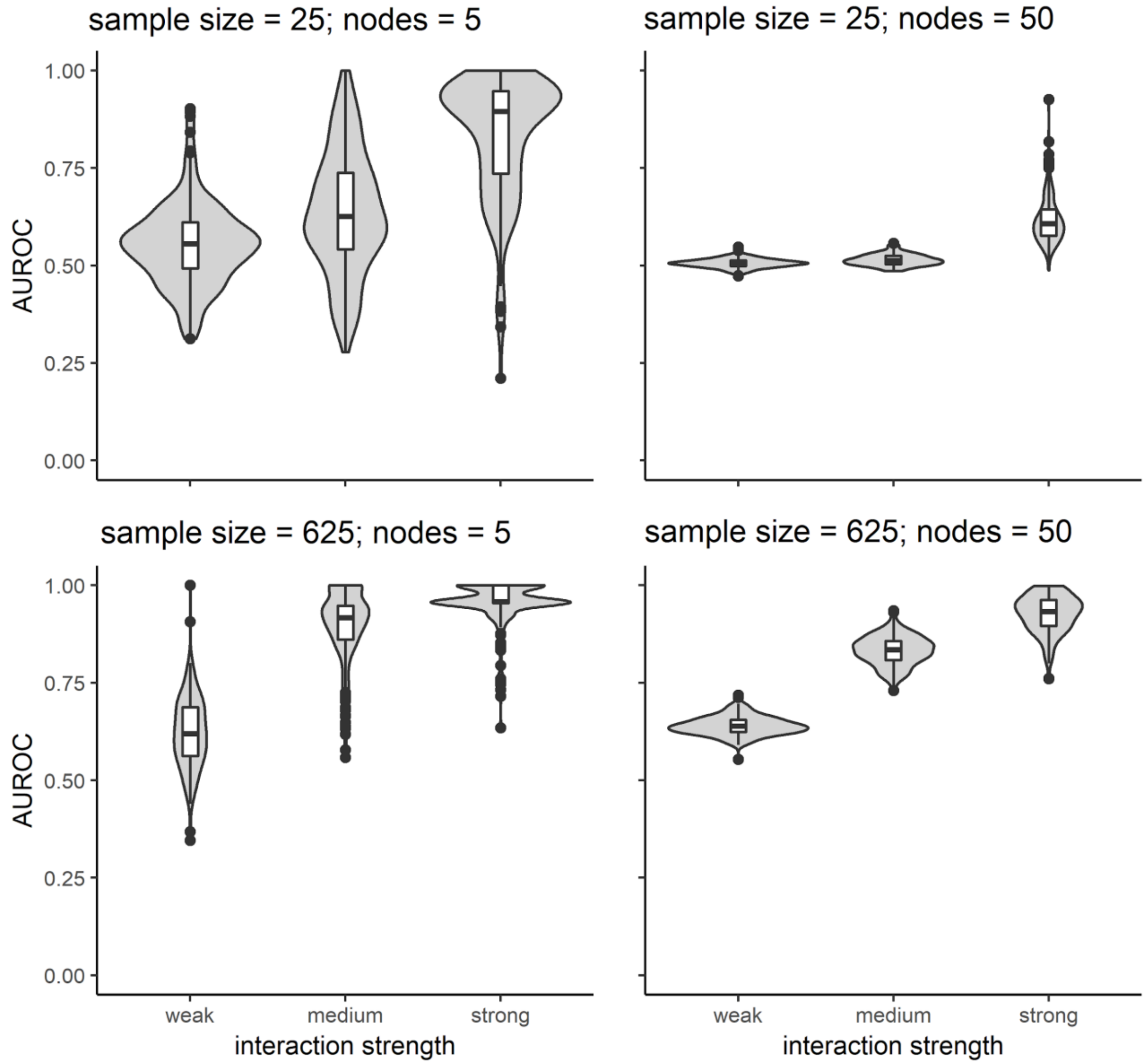


**Fig. S5.** Violin plots of the AUROC scores for the SBR method. The plot description is the same one found in Fig 3.

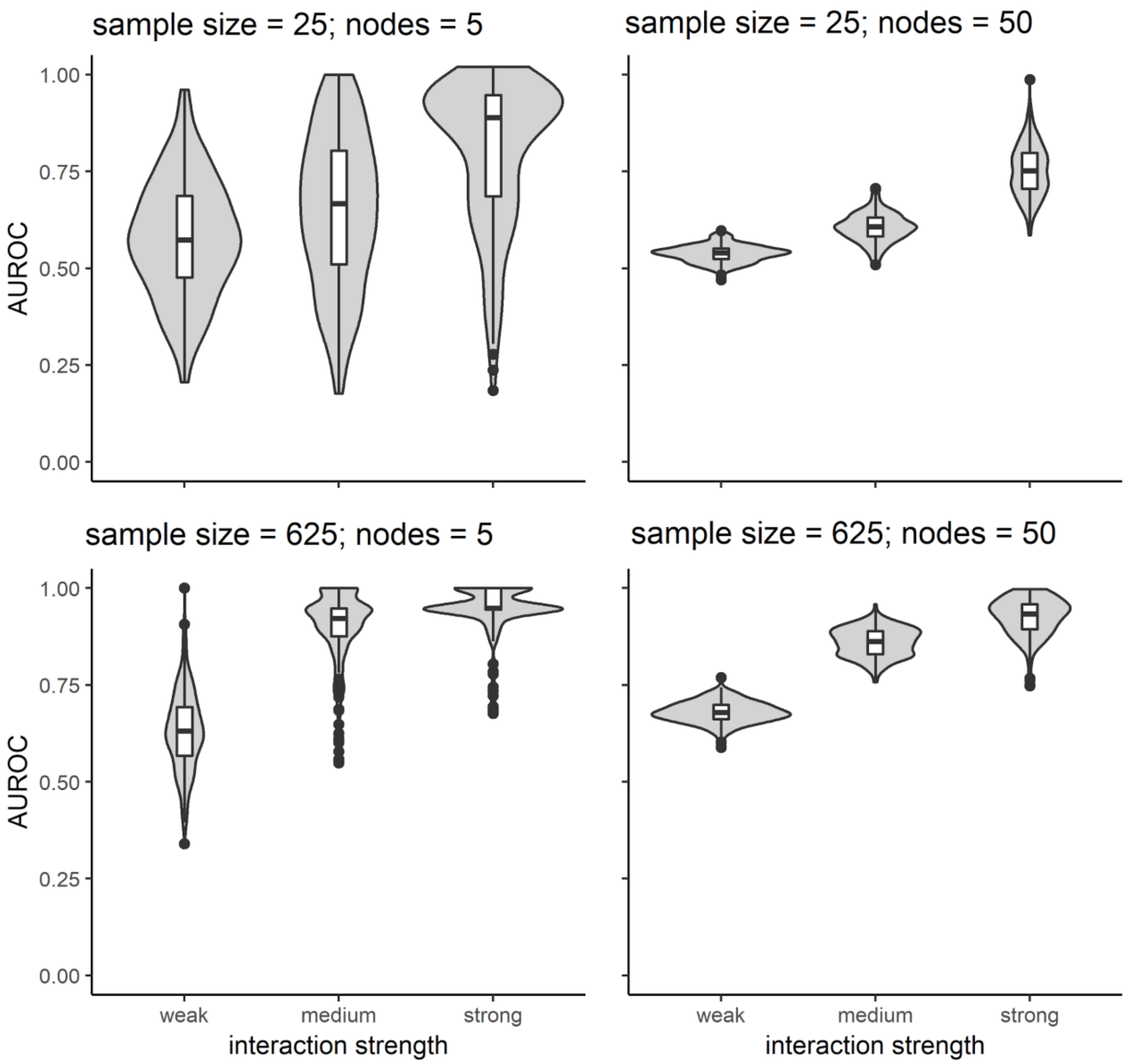


**Fig. S6.** Violin plots of the AUROC scores for the LASSO method. The plot description is the same one found in Fig 3.

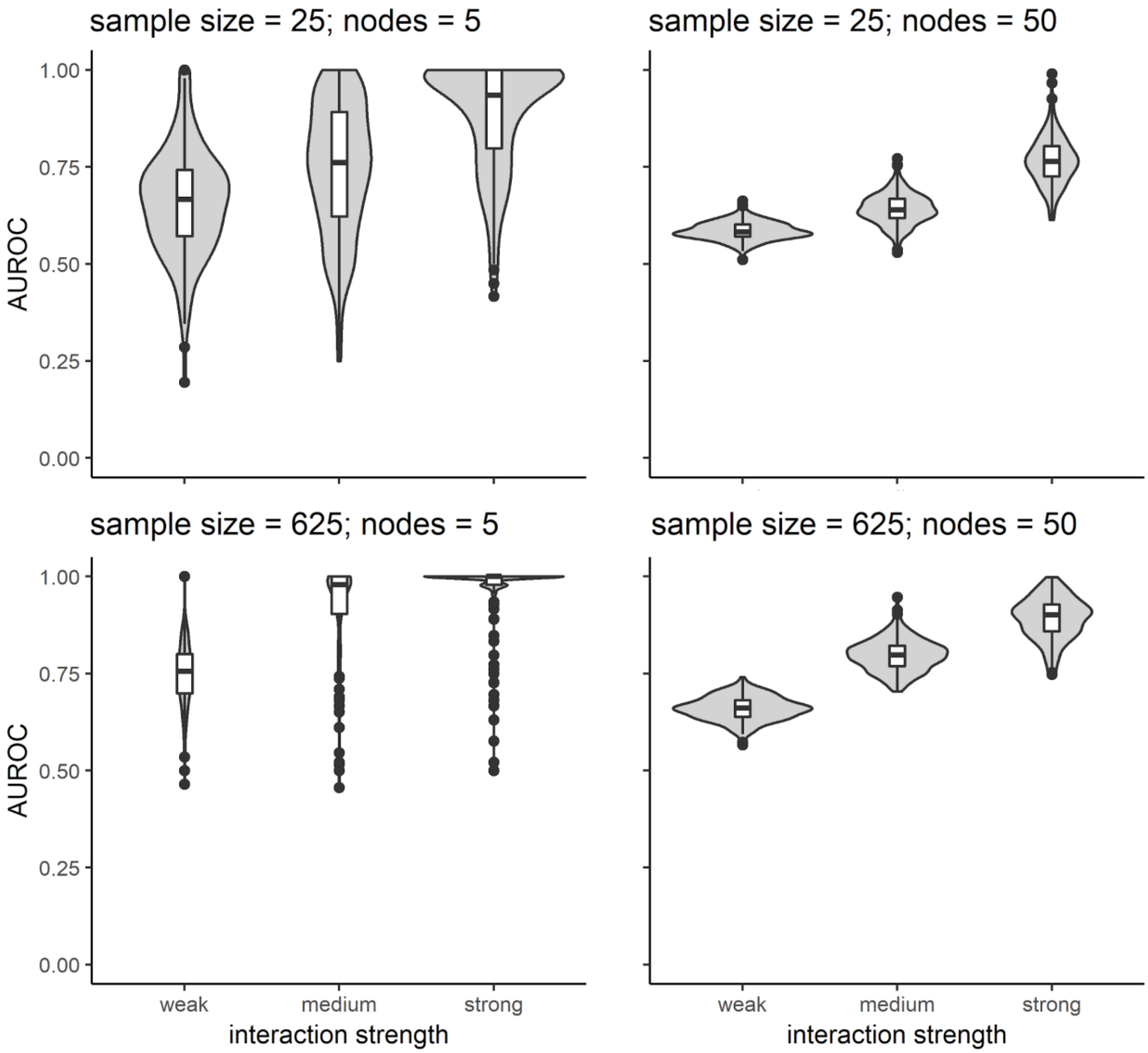


**Fig. S7.** Violin plots of the AUROC scores for the BN method. The plot description is the same one found in Fig 3.

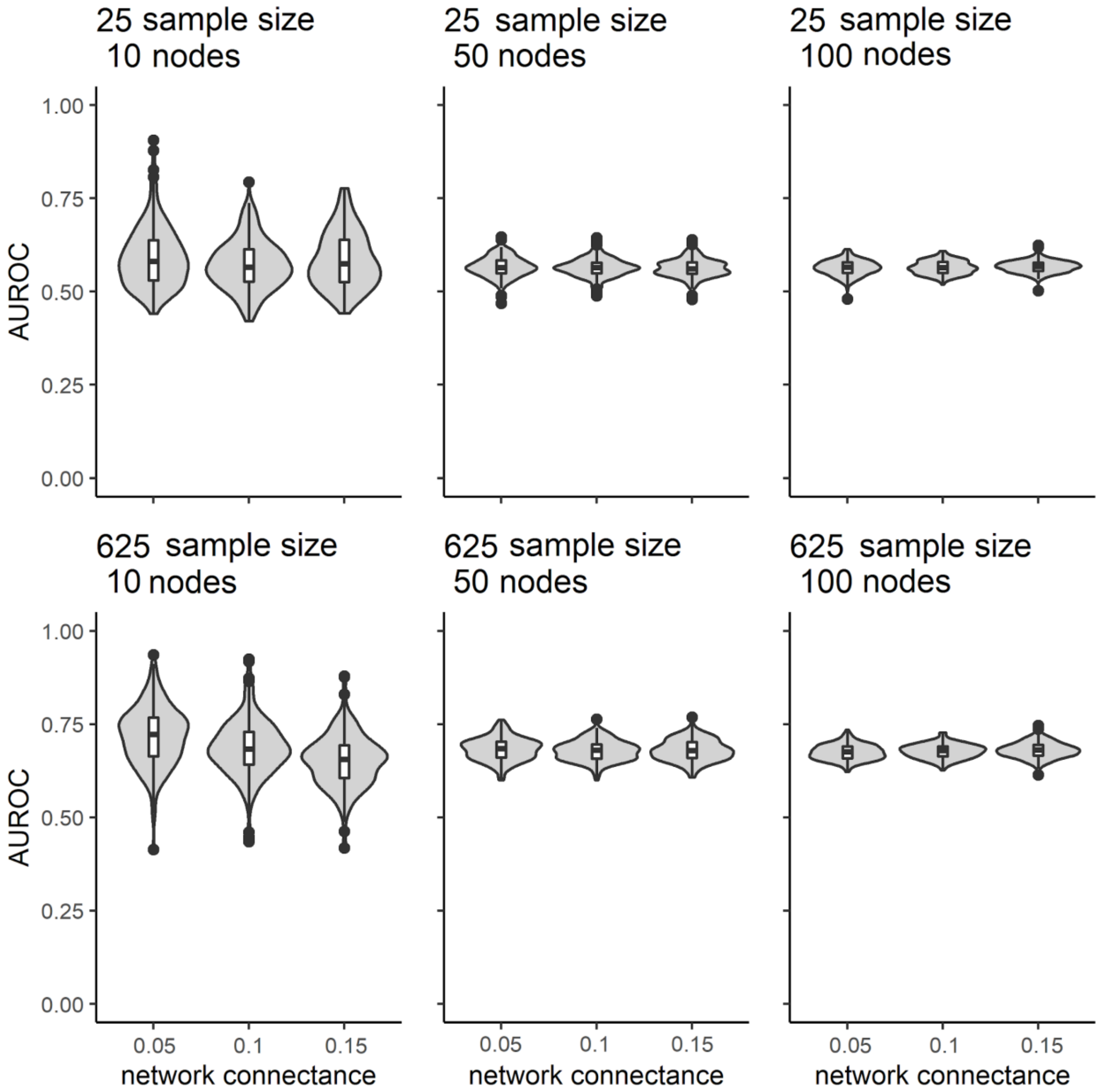


**Fig. S8.**

Violin plots of the AUROC scores for the GGM method with weak interaction strength and different network connectivity values. The plot description is the same one found in Fig 3.

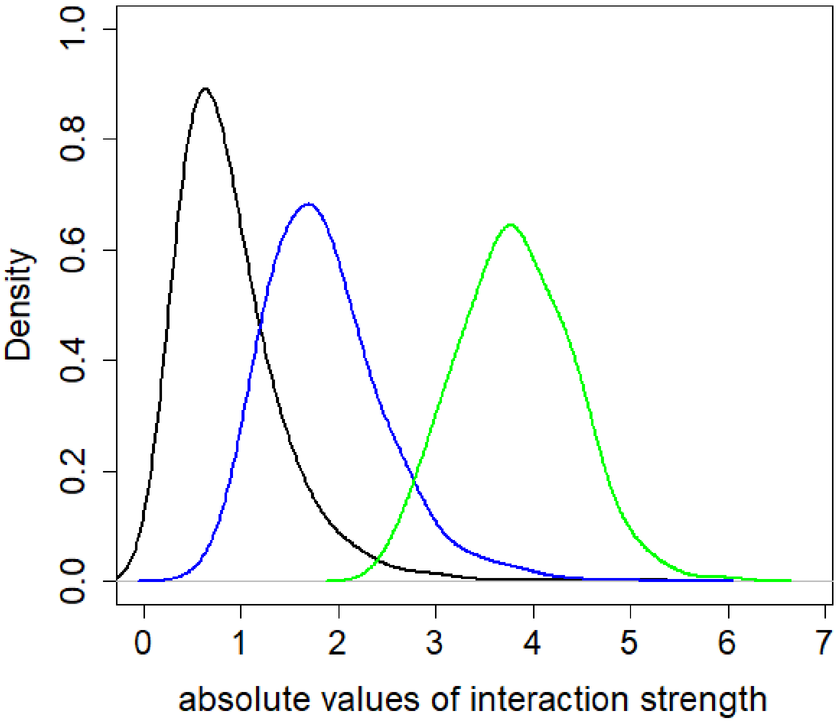


**Fig. S9.**

Density plots of absolute non-zero values of interaction strengths for weak interactions (black line), medium interactions (blue line) and strong interactions (green line). The density plots are smoothed distributions of the data points and the peaks of the plots are showing where there is a highest concentration of the data points (10 nodes and sample size = 625).

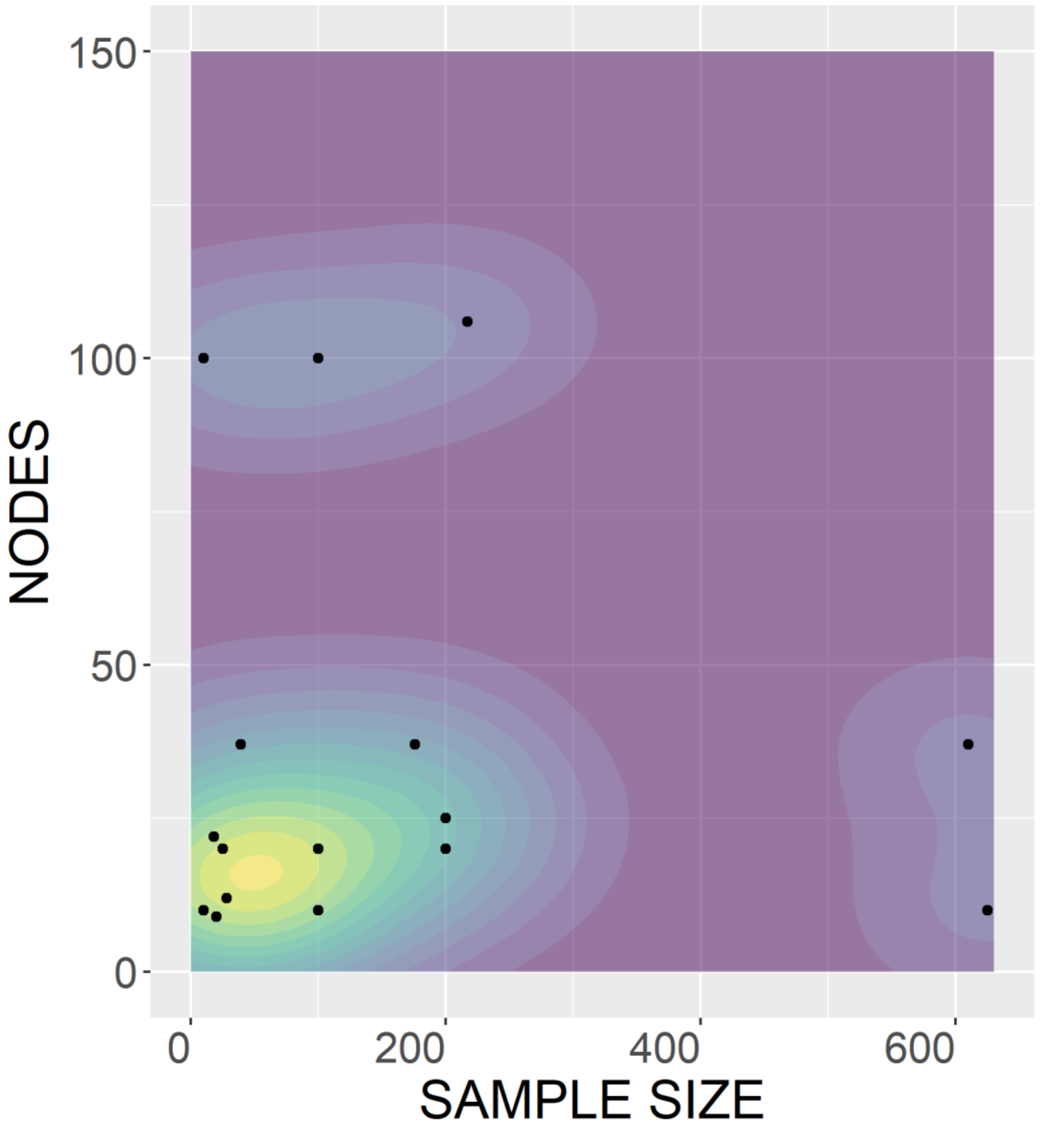


**Fig. S10.**

Parameter space (in terms of nodes-sample size) of sixteen published studies that used one or more of the considered network inference models. Yellow color shows high concentration of studies, green - moderate concentration of studies, light blue shows only few studies and purple color shows that no studies were found in that area among available online resources.
